# Automated model discovery for human cardiac tissue: Discovering the best model and parameters

**DOI:** 10.1101/2024.02.27.582427

**Authors:** Denisa Martonová, Mathias Peirlinck, Kevin Linka, Gerhard A. Holzapfel, Sigrid Leyendecker, Ellen Kuhl

**Affiliations:** Institute of Applied Dynamics, Friedrich-Alexander-Universität Erlangen-Nürnberg, Germany; Department of BioMechanical Engineering, Delft University of Technology, The Netherlands; Institute of Applied Mechanics, RWTH Aachen, Germany; Institute of Biomechanics, Graz University of Technology, Austria; Department of Structural Engineering, Norwegian University of Science and Technology, Norway; Department of Mechanical Engineering, Stanford University, California, United States

**Keywords:** automated model discovery, constitutive neural networks, constitutive modeling, machine learning, cardiac modeling

## Abstract

For more than half a century, scientists have developed mathematical models to understand the behavior of the human heart. Today, we have dozens of heart tissue models to choose from, but selecting the best model is limited to expert professionals, prone to user bias, and vulnerable to human error. Here we take the human out of the loop and automate the process of model discovery. Towards this goal, we establish a novel incompressible orthotropic constitutive neural network to simultaneously discover both, model and parameters, that best explain human cardiac tissue. Notably, our network features 32 individual terms, 8 isotropic and 24 anisotropic, and fully autonomously selects the best model, out of more than 4 billion possible combinations of terms. We demonstrate that we can successfully train the network with triaxial shear and biaxial extension tests and systematically sparsify the parameter vector with *L*_1_-regularization. Strikingly, we robustly discover a four-term model that features a quadratic term in the second invariant *I*_2_, and exponential quadratic terms in the fourth and eighth invariants *I*_4f_, *I*_4n_, and *I*_8fs_. Importantly, our discovered model is interpretable by design and has parameters with well-defined physical units. We show that it outperforms popular existing myocardium models and generalizes well, from homogeneous laboratory tests to heterogeneous whole heart simulations. This is made possible by a new universal material subroutine that directly takes the discovered network weights as input. Automating the process of model discovery has the potential to democratize cardiac modeling, broaden participation in scientific discovery, and accelerate the development of innovative treatments for cardiovascular disease.

Our source code, data, and examples are available at https://github.com/LivingMatterLab/CANN.

## 1 Motivation

Congenital heart defects, heart valve disease, and heart failure are just some of the many critical diseases of the heart that require medical intervention, for example in the form of corrective surgeries, valve repair or replacement, or cardiac assist devices [40]. In all these conditions and procedures, understanding the mechanics of cardiac tissue is crucial for diagnosis, treatment, and management, to optimize cardiac function and patient outcomes [36]. For more than half a century, scientists have developed mechanical models for human heart tissue [11, 20]. While the first models were purely *isotropic* [13], more sophisticated approaches soon acknowledged the importance of muscle fibers in the form of *transversely isotropic* [35] and *orthotropic* [27] models. All members of this first generation of models are *strain-based* Fung-type models [21], that embed a combination of directional strains into an exponential free energy function [22]. Unfortunately, this exponential mixed-term free energy function is not generally polyconvex [38, 65], and may violate the basic principles of thermodynamics [37].

Today, most popular models for heart muscle tissue are Holzapfel-type models [32] that use an *invariant-based* formulation of the free energy function [62] and can naturally incorporate tissue *incompressibility* and *orthotropy* in the fiber, sheet, and normal directions [33]. Invariant-based modeling of cardiac tissue has rapidly gained popularity [16, 23, 24, 30, 45, 46, 52] and is now widely used in many common finite element packages [1, 7]. Notably, the initial Holzapfel Ogden model was made up of four exponential quadratic terms in the first invariant *I*_1_, the fourth invariants *I*_4f_ and *I*_4s_, and the eighth invariant *I*_8fs_, with two parameters each; one with the unit of stiffness and the other unit-less [33]. While this initial four-term model performs well on simple shear tests of porcine heart tissue [14], it displays limitations when simultaneously fit to different loading modes [26]. Although the Holzapfel Ogden model is popular and widely used, a fair question to ask is, is this really the best possible model?

To answer this question, we abandon the common practice to a priori select a specific model, fit its parameters to data, and try to increase its goodness of fit [15, 59]. We also refrain from selectively adding or removing individual terms to incrementally improve an existing model [26]. Instead, we adopt the paradigm of *constitutive neural networks* [41] to autonomously discover the best model and parameters from a wide variety of possible terms [42]. The underlying idea is to generalize the Holzapfel Ogden model and design an orthotropic, perfectly incompressible constitutive neural network that takes the two isotropic invariants *I*_1_, *I*_2_ and the six anisotropic invariants *I*_4f_, *I*_4s_, *I*_4n_, *I*_8fs_, *I*_8fn_, *I*_8sn_ as input and approximates the free energy function as output. This network has two hidden layers: the first layer generates powers (º) and (º)^2^ of the invariants, and the second layer applies the identity (º) and exponential (exp(º)) to these powers [29, 44, 66]. This results in 8 *×* 2 *×* 2 = 32 terms, 48 model parameters, and 32^2^ = 4.294.967.296 possible models. To discover the best of these more than 4 billion models, we train our neural network with triaxial shear and biaxial extension data from human heart tissue [61]. In general, we expect the network to discover models with dense parameter vectors for which a subset of weights trains to zero [57]. Our intuition tells us that, the more non-zero weights we discover, the more complex the model, and the better the fit to the data [3, 50]. However, models with too many parameters and too many terms are difficult to interpret and generalize poorly to unseen data [9]. So a critical question to address is, how can we discover sparse models with only a few easy-to-understand terms?

A popular strategy to induce sparsity in a regression problem is *L*_p_-regularization [19]. *L*_p_-regularization adds the weighted *L*_p_-norm of the parameter vector to the loss function and induces sparsity for p-values equal to or smaller than one [28]. Here we induce sparsity using *L*_1_-regularization or lasso [67] by adding the weighted sum of the network weights to the loss function of our constitutive neural network [63]. This additional term allows us to fine-tune the number of non-zero parameters of our model; yet, at the expense of a reduced goodness-of-fit and at the cost of an additional hyperparameter, the penalty parameter *α* [48]. We can interpret this penalty parameter as a continuous switch between minimizing the network loss and minimizing the number of terms [18]. Clearly, the solution will be sensitive to this penalty parameter, but there are no obvious guidelines how to select the value of *α*. An important question in model discovery is therefore, how do we select the penalty parameter to best balance accuracy and sparsity?

The objective of this manuscript is to discover the model and parameters that best describe human heart tissue using the paradigm of constitutive neural networks, supplemented with *L*_1_-regularization. Towards this goal, we first summarize the basics of continuum mechanics in Section 2.1, and then integrate this knowledge into a new family of incompressible orthotropic constitutive neural networks in Section 2.2. In the Section 2.3, we briefly reiterate the deformation modes of triaxial shear and biaxial extension and summarize the experimental data we use to train our network. In Section 2.4, we introduce our finite element model to perform real heart simulation. In Section 3, we summarize our results of five overarching studies: (i) model discovery with triaxial shear, biaxial extension, and both data sets combined; (ii) model discovery with five different levels of regularization; (iii) model discovery with five different sets of initial conditions; (iv) specialization of our approach to three popular models; and (v) generalization of our results from homogeneous laboratory tests to heterogeneous real heart simulations. In Section 4 we discuss our results, limitations, and future directions, and close with a brief conclusion.

## 2 Methods

### 2.1 Continuum model

To characterize the deformation of the sample, we introduce the deformation map φ as the mapping of material points ***X*** in the undeformed configuration to points ***x*** = **φ** (***X***) in the deformed configuration [4, 32]. The gradient of the deformation map **φ** with respect to the undeformed coordinates ***X*** defines the deformation gradient ***F*** with its determinant *J*, and its right and left multiplications with its transpose ***F***^t^ define the left Cauchy-Green deformation tensor ***b***,

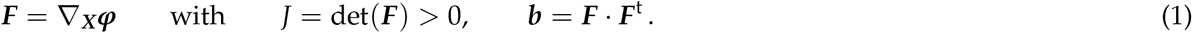

We assume that myocardial tissue is orthotropic, with three pronounced directions, ***f*** _0_, ***s***_0_, ***n***_0_, associated with the fiber, sheet, and normal directions in the reference configuration, where all three vectors are unit vectors, || ***f*** _0_ || = 1, || ***s***_0_ || = 1, || ***n***_0_ || = 1. We characterize the deformation in terms of nine invariants [47, 62] three standard isotropic invariants *I*_1_, *I*_2_, *I*_3_, three anisotropic invariants associated with the stretches squared, *I*_4f_, *I*_4s_, *I*_4n_, and three coupling invariants, *I*_8fs_, *I*_8fn_, *I*_8sn_,

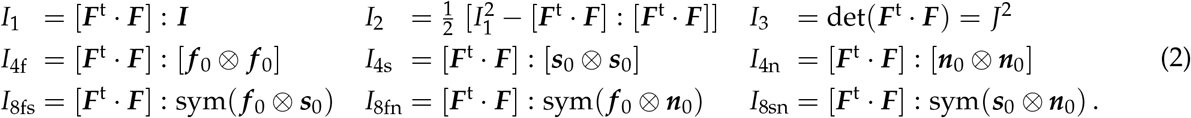

For convenience, we can reformulate the nine invariants in terms of the left Cauchy-Green deformation tensor ***b*** and the fiber, sheet, and normal directions in the deformed configuration, ***f*** = ***F*** *·* ***f*** _0_, ***s*** = ***F*** *·* ***s***_0_, ***n*** = ***F*** *·* ***n***_0_,

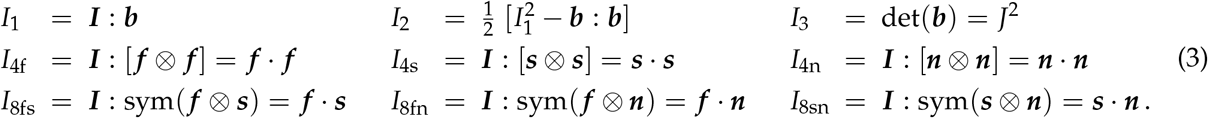

We assume that the tissue is perfectly incompressible, such that the third invariant remains constant and equal to one, *I*_3_ = *J*^2^ = 1. We then introduce the free energy function *ψ* as a function of the remaining eight invariants, *I*_1_, *I*_2_, *I*_4f_, *I*_4s_, *I*_4n_, *I*_8fs_, *I*_8fn_, *I*_8sn_,

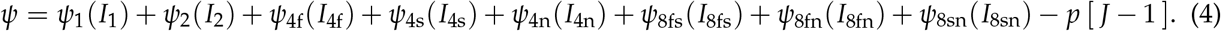

The last term, *p* [*J−* 1], enforces incompressibility and the pressure *p* acts as a Lagrange multiplier. For simplicity, we assume that all eight terms of the free energy function are uncoupled and none of the invariants directly influences the other seven. At this point, traditional constitutive modeling approaches assume a specific form of the free energy function *ψ*, and fit its parameters to data. Here, instead, we seek to *discover* the best free energy function *ψ* and the best parameters **w** = {*w*_i,j_} that explain our experimental data.

### 2.2 Neural network model

To automate the process of model discovery, we adopt the concept of constitutive neural networks [41]. Figure 1 illustrates the custom-designed architecture of our orthotropic, perfectly incompressible constitutive neural network with two hidden layers and 32 nodes that takes the eight invariants *I*_1_, *I*_2_, *I*_4f_, *I*_4s_, *I*_4n_, *I*_8fs_, *I*_8fn_, *I*_8sn_ as input and approximates the free energy ψ as output. The first layer generates powers (º) and (º)^2^ of the network input and the second layer applies the identity (º) and the exponential function (exp(º)) to these powers. This results in the following explicit representation of the free energy function ψ,

**Figure 1:**
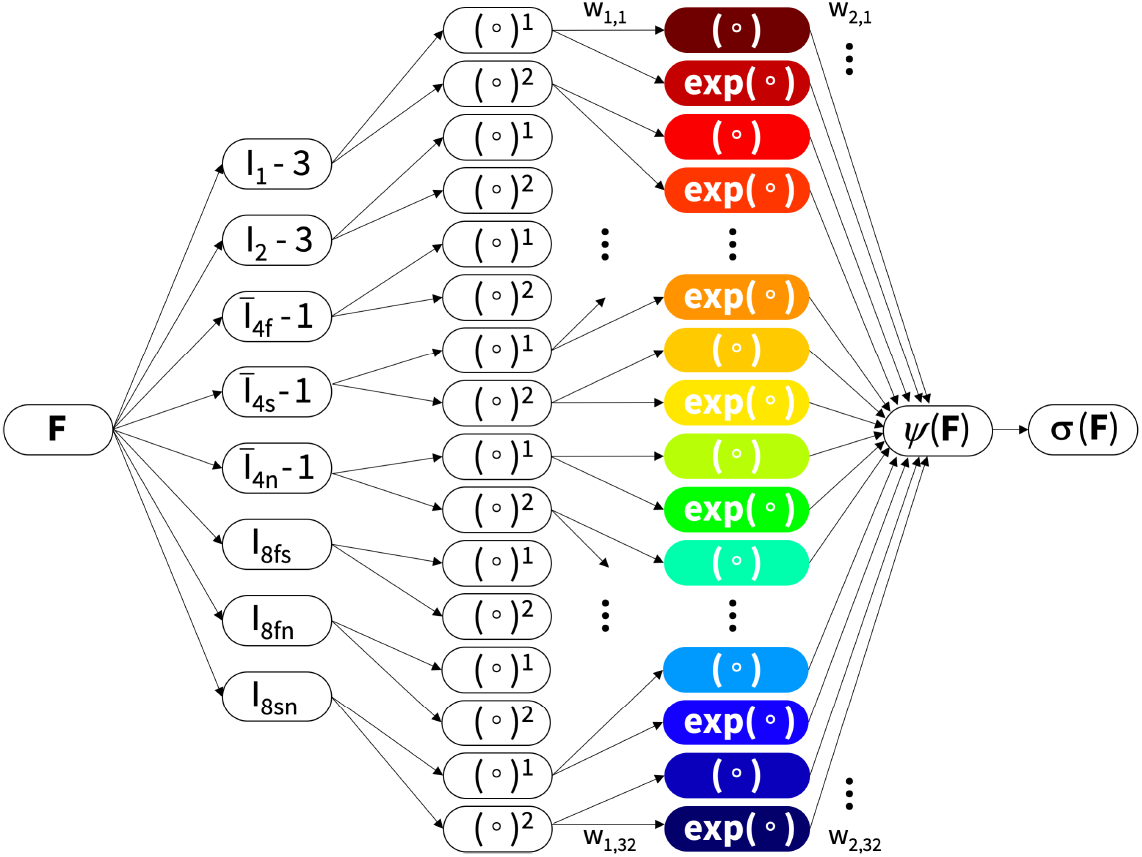
Constitutive neural network. Orthotropic, perfectly incompressible, feed forward constitutive neural network with two hidden layers to approximate the single scalar-valued free energy function, *ψ* (*I*_1_, *I*_2_, *I*_4f_, *I*_4s_, *I*_4n_, *I*_8fs_, *I*_8fn_, *I*_8sn_), as a function of eight invariants of the left Cauchy-Green deformation tensor ***b*** using 32 terms. The first layer generates powers (º) and (º)^2^ of the eight invariants and the second layer applies the identity (º) and exponential (exp(º)) to these powers. The network is not fully connected by design to satisfy the condition of polyconvexity a priori.

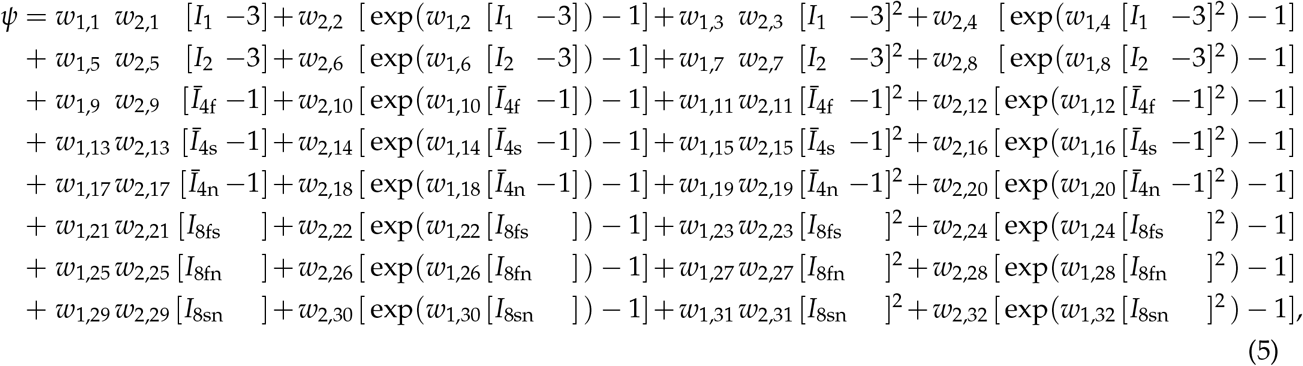

corrected by the pressure term *ψ* = *ψ − p* [*J −* 1]. For the isotropic invariants, *I*_1_, *I*_2_, the free energy function explicitly corrects for their values of three in the undeformed configuration using [*I*_1_ *−* 3], [*I*_2_ *−* 3]. For the anisotropic fourths invariants, *I*_4f_, *I*_4s_, *I*_4n_, the free energy function is only activated for tensile stretches [33], *Ī*_4f_ = max {*I*_4f_, 1}, *Ī*_4s_ = max {*I*_4s_, 1}, *Ī*_4n_ = max {*I*_4n_, 1}, and explicitly corrects for their values of one in the undeformed configuration using [*Ī*_4f_ *−* 1], [*Ī*_4s_ *−* 1], [*Ī*_4n_ *−* 1]. For the anisotropic eights invariants, *I*_8fs_, *I*_8fn_, *I*_8sn_, the values are zero in the undeformed configuration and can be used as is. We note, though, that the eighth invariants depend on the signs of the fiber, sheet, and normal directions and are therefore not strictly invariant [33]. From the free energy function *ψ* in Eq. (5), we can derive the Cauchy stress using standard arguments of thermodynamics,

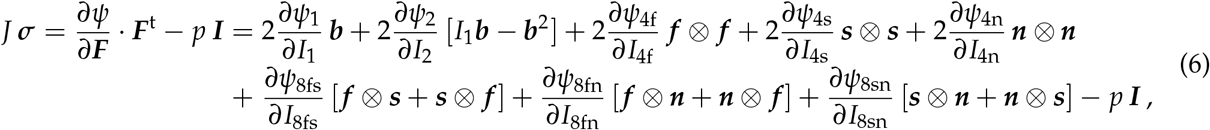

where the derivatives of the free energy with respect to the eight invariants take the following form,

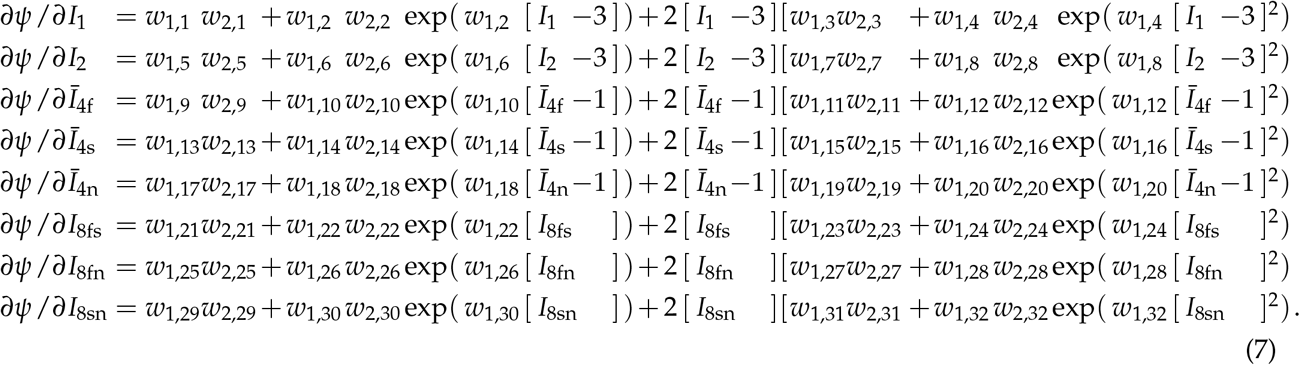

During training, the network learns the network weights **w** = {*w*_i,j_} which, by design, translate into physically meaningful model parameters that we enforce to always remain non-negative, **w** ≥ **0**. All weights of the first layer *w*_1,j_ are unit-less parameters, and all weights of the second layer *w*_2,j_ are stifnesslike parameters with the unit kilopascal. We note that all odd weights of the first layer are redundant. Without loss of generality, we can set them equal to one and reduce the total number of trainable weights from 64 to 48. We learn these network weights by minimizing a loss function *L* that penalizes the error between model and data. Here we use the mean squared error, the *L*_2_-norm of the difference between the model ***σ*** (***F***_*i*_, **w**) and the data 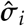, where *i*, 舰, *n*_data_ denotes the number of data points, divided by the total number of data points *n*_data_. To fine tune the number of weights in the model, we add a regularization term in the *L*_1_-norm of the *i*, 舰, *n*_weights_ weights, *α* || **w** ||_1_, to the loss function,

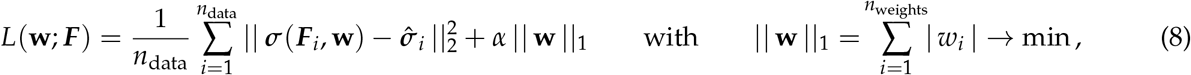

For penalty parameters *α*= 0, we recover the classical non-regularized constitutive neural network that simply minimizes the error between model and data. For penalty parameters *α* > *0, netw*ork training becomes an intricate balance between minimizing the error between model and data, and reducing the number of weights. Training some weights exactly to zero induces sparsity, effectively reduces model complexity, and improves interpretability [48]. To minimize the loss function in Eq. (8), we adopt the widely used ADAM optimizer, a robust adaptive algorithm for gradient-based first-order optimization, supplemented with an early stopping criterion for no accuracy change.

### 2.3 Triaxial shear and biaxial extension

The final step is to specify the eight invariants and the components of the Cauchy stress for our train and test data. We train our constitutive neural network with data from triaxial shear and biaxial extension tests on human myocardial tissue [61] and assume that the myocardium is orthotropic and perfectly incompressible. The *triaxial shear tests* used cubical specimens of 4 × 4 × 4 mm^3^, sheared in the fiber, sheet, and normal directions, resulting in six data sets of shear strain vs. shear stress pairs [33]. The *biaxial extension tests* used square specimens of 25 × 25 × 2.3 mm^3^, stretched in the fiber and normal directions at stretch ratios of 1:1, 1:0.75, 0.75:1, 1:0.5, 0.5:1 resulting in five times two data sets of stretch vs. normal stress pairs [61]. Table 1 summarizes the digitized data of the six triaxial shear tests and five biaxial extension tests [61]. For both experiments, the deformation gradient ***F*** and the Cauchy stress ***σ*** take the following format, where the subscripts, f, s, n, refer to the fiber, sheet, and normal directions, and the stress tensor is symmetric, *σ*_fs_ = *σ*_sf_, *σ*_fn_ = *σ*_nf_, *σ*_ns_ = *σ*_sn_, according to the balance of angular momentum,

**Table 1:**
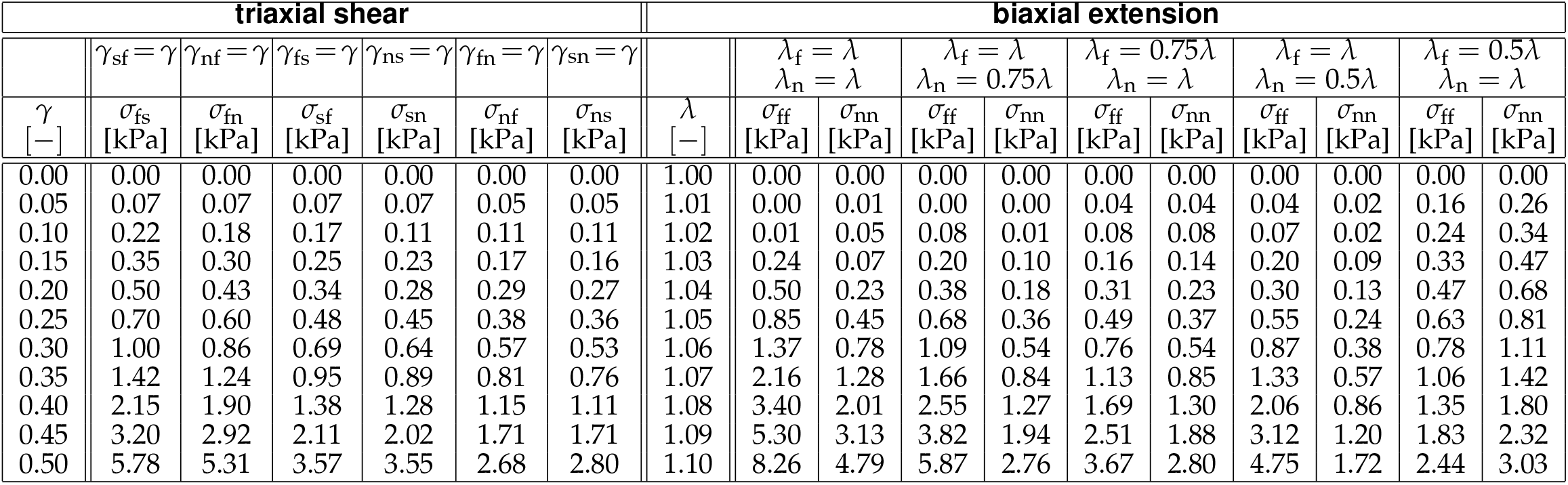
Triaxial shear and biaxial extension data for human myocardium. Human myocardial samples are sheared in six directions and stretched in two orthogonal directions at five different stretch ratios [61]. The indices f, s, n denote the fiber, sheet, and normal directions, where f and n are associated with the mean fiber and cross fiber directions MFD and CFD.

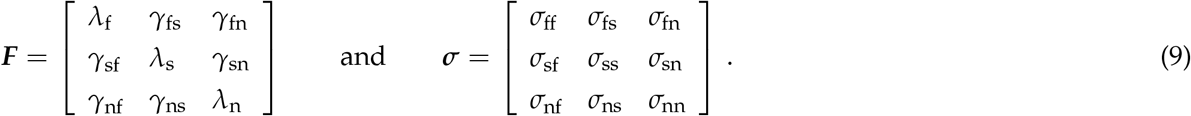

For the six *triaxial shear tests*, all three stretches remain constant to one, *λ*_f_ = *λ*_s_ = *λ*_n_ *≡* 1, and all but one shear term remain constant to zero, *γ*_fs_ = *γ*_sf_ = *γ*_nf_ = *γ*_fn_ = *γ*_sn_ = *γ*_ns_ *≡* 0. Each shear test is associated with one non-zero shear strain, and results in one non-zero pair of shear stresses, *σ*_fs_ = *σ*_sf_, *σ*_fn_ = *σ*_nf_, *σ*_ns_ = *σ*_sn_,

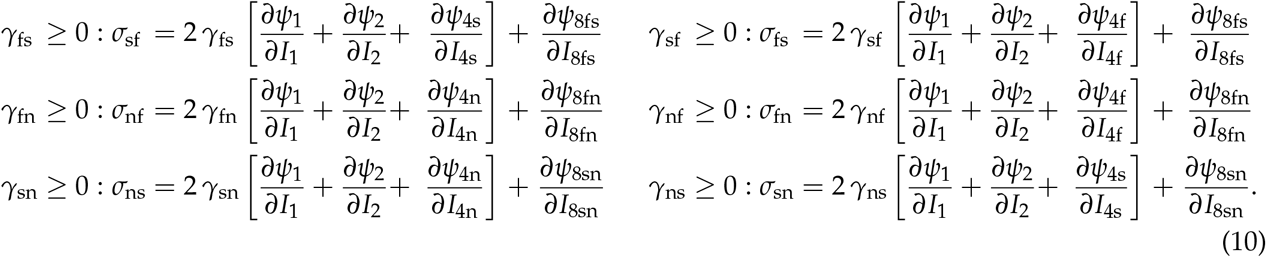

For the five *biaxial extension tests*, we vary the ratio of the fiber and normal stretches, *λ*_f_ and *λ*_n_, determine the sheet stretch, *λ*_s_ = 1/[*λ*_f_*λ*_n_] from the incompressibility condition, and keep all shear strains constant to zero, *γ*_fs_ = *γ*_sf_ = *γ*_nf_ = *γ*_fn_ = *γ*_sn_ = *γ*_ns_ *≡* 0. We derive the hydrostatic pressure *p* from the zero-normal-stress condition, *σ*_ss_ = 0, using Eq. (6), which results in the following non-zero normal stresses,

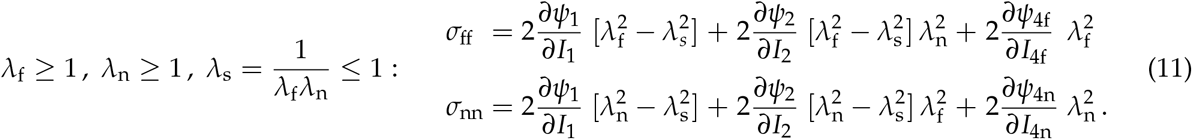

The definitions of the shear and normal stresses in Eqs. (10) and (11) use the derivatives of the free energy with respect to the eight invariants in Eq. (7), which are parameterized in terms of the network weights that we learn by minimizing the loss function in Eq. (8).

### 2.4 Real heart simulations

To probe the performance of our discovered models within a realistic real heart simulation, we incorporate our newly discovered models in the finite element analysis software solver Abaqus [1], and predict the stress state of the left and right ventricular wall during diastolic filling. We create a finite element model of the left and right ventricles from high-resolution magnetic resonance images of a healthy 44-year-old Caucasian male with a height of 178cm and weight of 70kg [51, 54]. We discretize the myocardial wall using 99,286 quadratic tetrahedral elements, with a total of 462,498 degrees of freedom. We incorporate the tissue microstructure through a helically wrapping fiber architecture in terms of 99,286 local fiber, sheet, and normal directions, ***f*** _0_, ***s***_0_, ***n***_0_. We compute these local microstructural orientations by solving a Laplace-Dirichlet problem across our computational domain, and assuming a transmural fiber variation from +60° to -60 ° from the endocardial to the epicardial wall [72]. To fix the ventricles in space, we apply homogeneous Dirichlet boundary conditions at the mitral, aortic, tricupid, and pulmonary valve annuli [53]. To translate our newly discovered orthotropic material models for myocardial tissue into a finite element analysis environment, we adopt our new *universal material model subroutine* [55, 56]. Here, each free energy contribution from our constitutive neural network in Figure 1 translates into one line of the parameter table that describes the term’s invariant, first-layer activation function, second-layer activation function, and weights *w*_1,*•*_ [-] and *w*_2,*•*_ [kPa].

## 3 Results

We successfully trained our constitutive neural network from Figure 1, first with the six triaxial shear tests, then with the five biaxial extension tests, and finally with all eleven tests simultaneously [61]. When only using the triaxial shear tests for training, we use the biaxial extension tests for testing, and vice versa. For all cases, the loss function converges robustly within 30,000 epochs, with an early stopping criterion for no accuracy change, see Table 2, bottom row. With a batch size of 32, each training run takes between one and eight hours using Google Colab. The computational time varies between individual and simultaneous training, and depends on the amount of training data and the regularization level. For each training set, in every direction, we compare the experimentally reported stress-shear and stress-stretch data to the discovered stress-shear and stress-stretch model. We report the goodness of fit in terms of the correlation coefficient, R^2^, and the root-mean-square error, rms, for both training and testing. We also report both the means of both metrics across all 16 testing modes, six for triaxial shear and five for biaxial extension in each of the two directions.

**Table 2:** Model discovery for human myocardium trained on triaxial shear and biaxial extension test for varying regularization levels. Discovered material parameters for simultaneous training with six shear and five biaxial tests for varying regularization levels *α*; mean goodness of fit R^2^ and root mean squared error rms; and number of epochs towards convergence.

| network weights<br>$w_{1,\bullet}$ [-], $w_{2,\bullet}$ [kPa] | model term<br>[-] | regularization level | | | | |
| --- | --- | --- | --- | --- | --- | --- |
| | | $\alpha = 0.0$ | $\alpha = 0.001$ | $\alpha = 0.01$ | $\alpha = 0.1$ | $\alpha = 1.0$ |
| $w_{1,5} \cdot w_{2,5}$ | $[I_2 - 3]$ | 0.034 | – | – | – | – |
| $w_{1,6}, w_{2,6}$ | $\exp([I_2 - 3])$ | 0.037, 8.454 | – | – | – | 2.100, 0.926 |
| $w_{1,7} \cdot w_{2,7}$ | $[I_2 - 3]^2$ | 4.648 | – | 5.162 | 5.978 | – |
| $w_{1,8}, w_{2,8}$ | $\exp([I_2 - 3]^2)$ | – | 0.368, 13.291 | – | – | – |
| $w_{1,12}, w_{2,12}$ | $\exp([\bar{I}_{4f} - 1]^2)$ | 24.328, 0.063 | 22.530, 0.073 | 21.151, 0.081 | 6.626, 0.389 | – |
| $w_{1,16}, w_{2,16}$ | $\exp([\bar{I}_{4s} - 1]^2)$ | 19.489, 0.028 | – | – | – | – |
| $w_{1,20}, w_{2,20}$ | $\exp([\bar{I}_{4n} - 1]^2)$ | 11.761, 0.095 | 9.618, 0.126 | 4.371, 0.315 | 1.211, 1.079 | – |
| $w_{1,21} \cdot w_{2,21}$ | $[I_{8fs}]$ | 0.100 | 0.108 | – | – | – |
| $w_{1,24}, w_{2,24}$ | $\exp([I_{8fs}]^2)$ | – | 5.478, 0.011 | 0.508, 0.486 | – | – |
| $w_{1,29} \cdot w_{2,29}$ | $[I_{8sn}]$ | 0.004 | – | – | – | – |
| $R^2$ | | 0.896 | 0.892 | 0.894 | 0.872 | 0.640 |
| rms |  | 0.409 | 0.423 | 0.426 | 0.460 | 0.644 |
| epochs |  | 30 000 | 27 900 | 26 400 | 12 700 | 4 400 |

### Rich training data are critical to discover generalizable models

In the first set of examples, we train the neural network with three different train-test scenarios: training with triaxial shear and testing with biaxial extension; training with biaxial extension and training with triaxial shear; training with both triaxial shear and biaxial extension [61]. In this first set of examples, we do not apply any regularization. Figure 2, top, and Figure 3, bottom, summarize the results for *training* with the triaxial shear data and with the biaxial extension data only. In both cases, the network trains well with a goodness of fit above R^2^ = 0.989 for all shear tests and above R^2^ = 0.924 for all biaxial extension tests with maximum fiber stretches of *λ*_f_ = 1.10, first, second, and fourth column. The quality of training is compromised for smaller fiber stretches, with a goodness of fit as low as R^2^ = 0.712 for the smallest maximum fiber stretches of *λ*_f_ = 1.05, fifth column. Figure 2, bottom, and Figure 3, top, summarize the results for *testing* with the biaxial extension and triaxial shear data, when trained with the other data set. In both cases, the predictions are poor with a mean goodness of fit of R^2^ = 0.532 and rms = 0.558 for training with the triaxial shear data and R^2^ = *−* 2.462 and rms = 1.411 for training with the biaxial extension data. Figure 4 summarizes the results for *simultaneous training* with the triaxial shear and biaxial extension data combined. Notably, the overall goodness of fit improves significantly with values above R^2^ = 0.789 for shear and above R^2^ = 0.719 for biaxial extension, and mean values of R^2^ = 0.896 and rms = 0.409 across all eleven data sets. Interestingly, for simultaneous training with all eleven data sets combined, we robustly discover an *eight-term model*,

**Figure 2:**
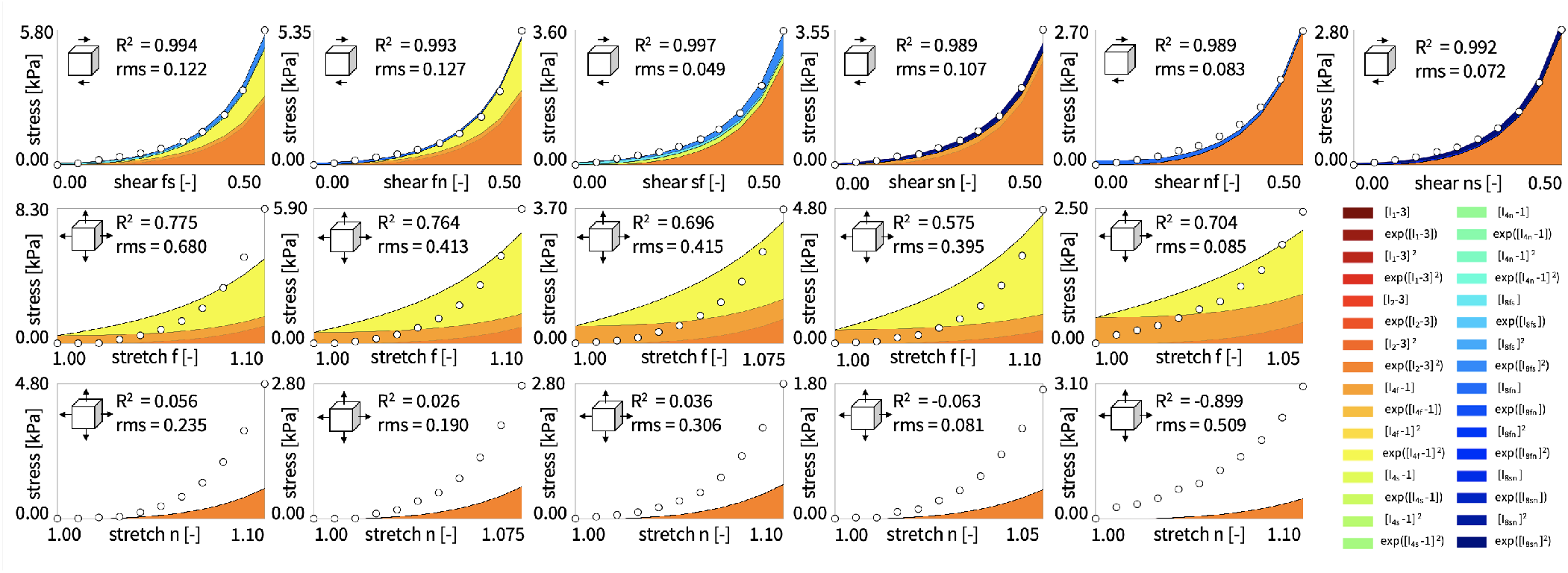
Model discovery for human myocardium trained on triaxial shear tests. Cauchy stress components as functions of shear strain during triaxial shear used for training, first row, and stretches during biaxial extension used for testing in fiber and normal directions, second and third rows, for the orthotropic, perfectly incompressible constitutive neural network with two hidden layers and 32 nodes from Figure 1. Dots illustrate the the experimental data [61] from Table 1; color-coded areas highlight the 32 contributions to the discovered stress function.

**Figure 3:**
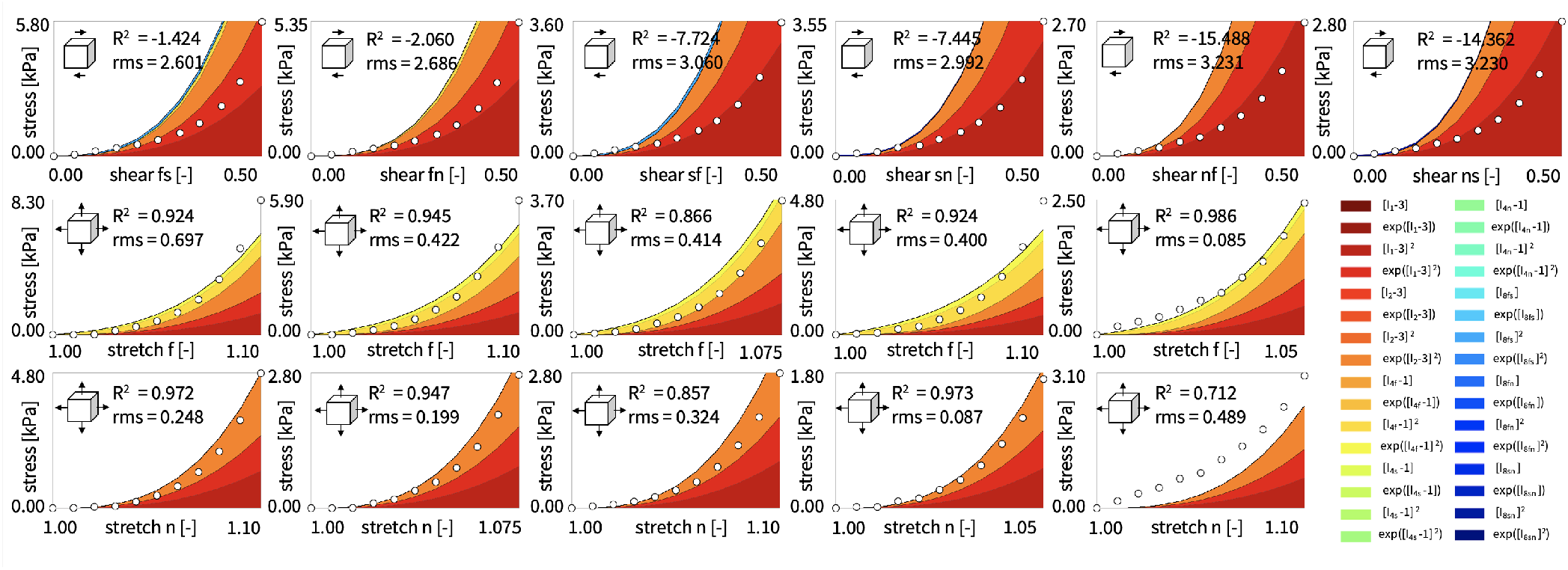
Model discovery for human myocardium trained on biaxial extension tests. Cauchy stress components as functions of stretches during biaxial extension in fiber and normal directions used for training, second and third rows, and shear strain during triaxial shear used for testing, first row, for the orthotropic, perfectly incompressible constitutive neural network with two hidden layers and 32 nodes from Figure 1. Dots illustrate the the experimental data [61] from Table 1; color-coded areas highlight the 32 contributions to the discovered stress function.

**Figure 4:**
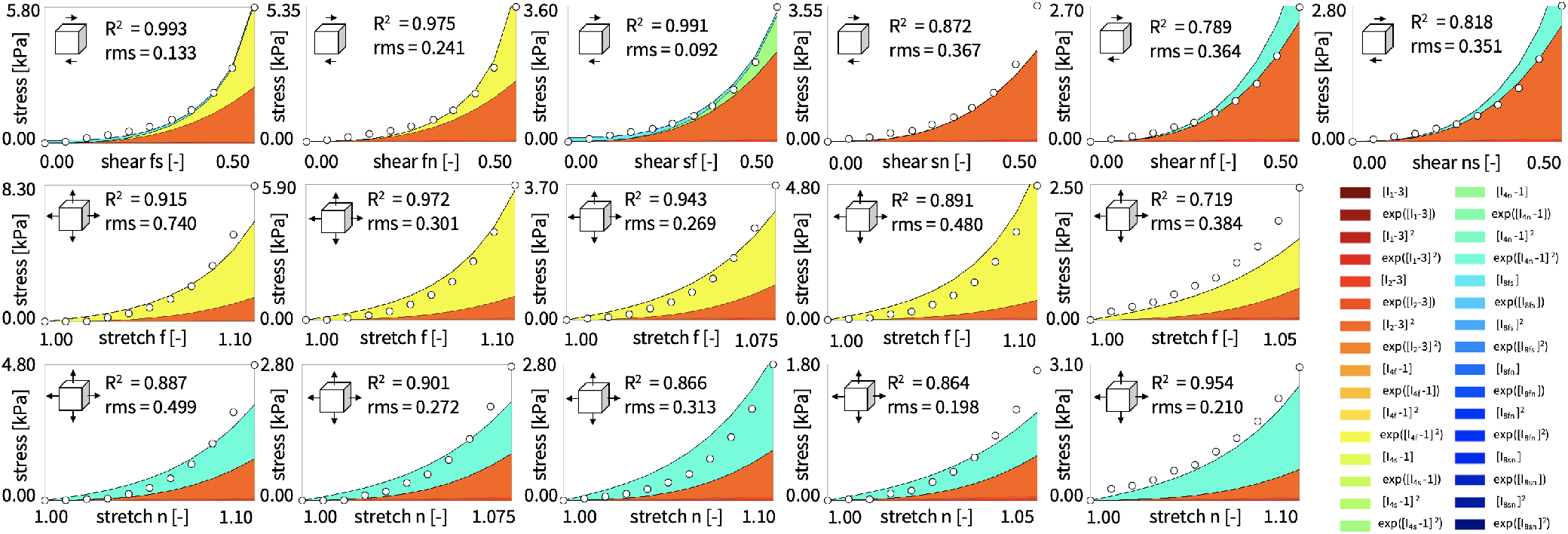
Model discovery for human myocardium trained on triaxial shear and biaxial extension tests. Cauchy stress components as functions of shear strain during triaxial shear, first row, and stretches during biaxial extension in fiber and normal directions, second and third rows, all used for training of the orthotropic, perfectly incompressible constitutive neural network with two hidden layers and 32 nodes from Figure 1. Dots illustrate the the experimental data [61] from Table 1; color-coded areas highlight the 32 contributions to the discovered stress function.

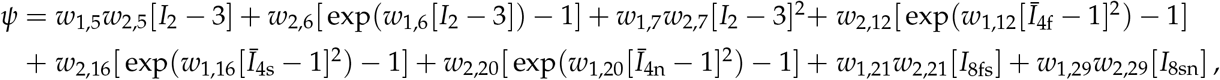

with linear, exponential linear, and quadratic terms in the second invariant *I*_2_, exponential quadratic terms in all fourth invariants **Ī**_4f_, **Ī**_4s_, **Ī**_4n_, and linear terms in two of the eighth invariants *I*_8fs_, *I*_8sn_. Strikingly, the discovered model does not contain a single term in the first invariant *I*_1_, and the isotropic response is entirely represented through the second invariant *I*_2_. Notably, all other 24 terms, including the classical neo Hooke term, naturally train to zero, even without any regularization. Table 2 summarizes the discovered non-zero weights of the discovered eight-term model. In total, the discovered model contains twelve parameters,

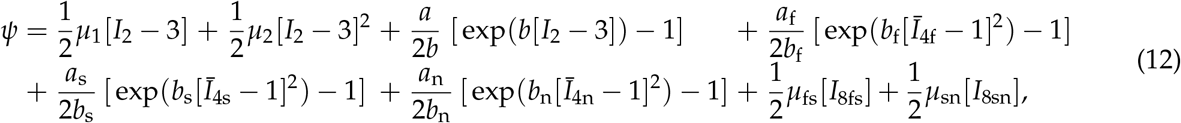

the eight stifness-like parameters *μ*_1_ = 2*w*_1,5_*w*_2,5_ = 0.068 kPa, *μ*_2_ = 2*w*_1,7_*w*_2,7_ = 9.296 kPa, *a* = 2*w*_1,6_*w*_2,6_ = 0.626 kPa, *a*_f_ = 2*w*_1,12_*w*_2,12_ = 3.065 kPa, *a*_s_ = 2*w*_1,16_*w*_2,16_ = 1.091 kPa, and *a*_n_ = 2*w*_1,20_*w*_2,20_ = 2.235 kPa, *a*_fs_ = 2*w*_1,21_*w*_2,21_ = 0.200 kPa, *a*_sn_ = 2*w*_1,29_*w*_2,29_ = 0.008 kPa, and the four exponents *b* = *w*_1,6_ = 0.037, *b*_f_ = *w*_1,12_ = 24.328, *b*_s_ = *w*_1,16_ = 19.489 and *b*_n_ = *w*_1,20_ = 11.761. When comparing the parameter values with the dominant colors in Figure 4, we conclude that the orange quadratic second invariant *I*_2_ term, the yellow exponential quadratic fourth invariant **Ī**_4f_ term, and the turquoise exponential quadratic fourth invariant **Ī**_4n_ term associated with the nodes 7, 12, 20 of our network in Figure 1 are the most relevant terms to represent the behavior of myocardial tissue. Taken together, these results suggest that, to discover the best model for myocardial tissue, it is critical to train the network on both triaxial shear and biaxial extension data simultaneously. From now on, we will only use both tests simultaneously and discover modes for all eleven data sets combined.

### Sparse regression promotes interpretable models

In the next set of examples, we explore the role of *L*_1_-regularization to induce sparsity in the discovered models [48]. We systematically increase the penalty parameter *α* in the loss function in Eq. (8), *α* = [0.0, 0.001, 0.01, 0.1, 1.0], and study its effect on the number of non-zero terms and the goodness of fit. Table 2 and Figure 5 summarize the resulting network weights and stress-shear and stress-stretch relations for the discovered models. The discovery converges robustly in all five cases, but requires progressively fewer epochs towards convergence, [30000, 27900, 26400, 12700, 4400]. The model without regularization, *α* = 0.0, is the eight-term model in Figure 4 in the previous section. This study confirms our general intuition that *L*_1_-regularization is an intricate balance between model sparsity and model accuracy, and that the penalty parameter *α* serves to fine-tune and down-select the number of relevant terms: As the penalty parameter increases, the network discovers progressively fewer non-zero terms, *n* = [8, 5, 4, 3, 1]. At the same time, the mean goodness of fit across all eleven data sets decreases, R^2^ = [0.896, 0.892, 0.894, 0.872, 0.640], and the mean root mean squared error increases, rms = [0.409, 0.423, 0.426, 0.460, 0.644]. For *α* = 0.01, we robustly discover a *four-term model*,

**Figure 5:**
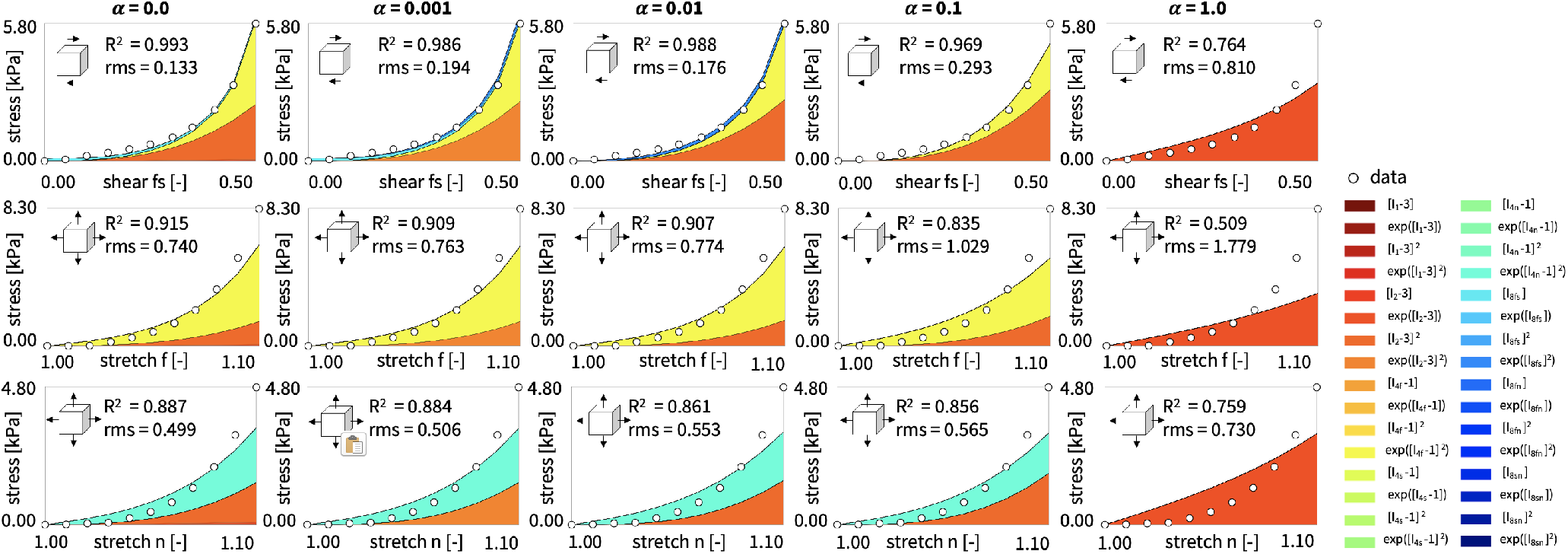
Model discovery for human myocardium trained on triaxial shear and biaxial extension test for varying regularization levels. Effect of penalty parameter *α* = { 0, 0.001, 0.01, 0.1, 1} for *L*_1_-regularization to induce sparsity. Cauchy stress components as functions of shear strain during triaxial shear, first row, and stretches during biaxial extension in fiber and normal directions, second and third rows, all used for training of the orthotropic, perfectly incompressible constitutive neural network with two hidden layers and 32 nodes from Figure 1. Dots illustrate the the experimental data [61] from Table 1; color-coded areas highlight the 32 contributions to the discovered stress function.

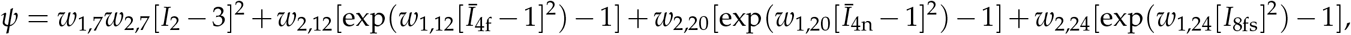

with a quadratic term in the second invariant *I*_2_, exponential quadratic terms in the fiber and normal fourth invariants **Ī**_4f_ and **Ī**_4n_, and an exponential quadratic term in the fiber-sheet eighth invariant *I*_8fs_. Increasing the penalty parameter to *α* = 0.1 results in a *three-term model*, that is a special case of this four-term model, independent of the eighth invariant,

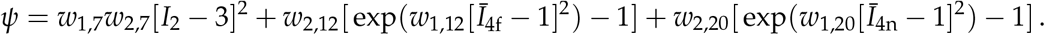

Clearly, the model with the largest penalty parameter of *α* = 1.0, the *one-term model*,

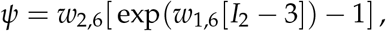

with an exponential linear second invariant *I*_2_ term, is incapable of capturing anisotropy and is not well suited to represent myocardial tissue. Its goodness of fit is significantly lower than that of all other models, suggesting that a penalty parameter of *α* = 1.0 over-regularizes model discovery and is simply too large to provide a reasonable fit. Figure 5 illustrates selected shear and biaxial tests for the five different regularization levels. Interestingly, the trend towards three relevant terms is clearly visible when comparing the the *α* = 0.0, *α* = 0.01, and *α* = 0.1 regularization, and their discovered eight-term, four-term, and three-term models that all contain the same dominant quadratic second invariant *I*_2_ term in orange, exponential quadratic fourth invariant **Ī**_4f_ term in yellow, and exponential quadratic fourth invariant **Ī**_4n_ term in turquoise, with only minor modifications when adding additional terms. The striking dominance of the orange, yellow, and turquoise terms, which already stood out prominently in the non-regularized eight-term model of Eq. (12), suggests that the best models to characterize the most important mechanical features of myocardial tissue are the *four-term model*,

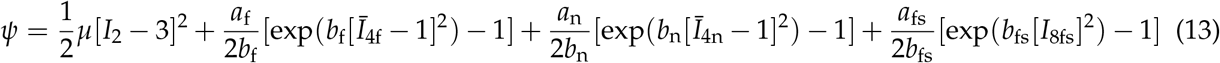

with seven material parameters, the four stiffness-like parameters *μ* = 2*w*_1,7_*w*_2,7_ = 10.324 kPa, *a*_f_ = 2*w*_1,12_*w*_2,12_ = 3.427 kPa, *a*_n_ = 2*w*_1,20_*w*_2,20_ = 2.754 kPa, and *a*_fs_ = 2*w*_1,24_*w*_2,24_ = 0.494 kPa, and the three exponents *b*_f_ = *w*_1,12_ = 21.151 *b*_n_ = *w*_1,20_ = 4.371, and *b*_fs_ = *w*_1,24_ = 0.508, and the *three-term model*,

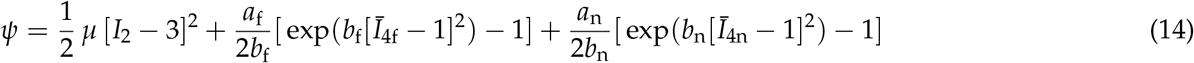

with the five material parameters, the three stiffness-like parameters *μ* = 2*w*_1,7_*w*_2,7_ = 11.956 kPa, *a*_f_ = 2*w*_1,12_*w*_2,12_ = 5.155 kPa, and *a*_n_ = 2*w*_1,20_*w*_2,20_ = 2.613 kPa, and the two exponents *b*_f_ = *w*_1,12_ = 6.626 and *b*_n_ = *w*_1,20_ = 1.211, according to Table 2.

### Model discovery is robust and relatively insensitive to the initial conditions

To demonstrate the robustness of our model discovery, independent of the initial conditions for our model parameters, we perform a series of random initializations and compare the discovered models. Table 3 confirms that, independent of the initial guess, our constitutive neural network robustly discovers the same best four-term model and the same best seven parameters, with only minor deviations. For brevity, we only show the results for a regularization level of *α* = 0.01, but emphasize that all other discovered models in Table 2 are equally robust to their initialization. For the displayed regularization level, for all five models, the invariants *I*_2_, *I*_4f_, *I*_4n_, *I*_8fs_ contribute quadratically to the free energy. The mean goodness of fit is largest, R^2^ = 0.894, and the mean error is smallest, rms = 0.426, for the first, second, and fourth initializations, which all contain the same four terms in the quadratic second invariant *I*_2_, exponential quadratic fourth invariants **Ī**_4f_ and **Ī**_4n_, and exponential quadratic eighth invariants *I*_8fs_, the orange, yellow, turquoise, and blue terms in Figure 5. Taken together, this example confirms that, although the loss function of our minimization problem in Eq. (8) is non-convex, our model discovery is robust and consistently discovers similar models with similar parameter values.

**Table 3:** Model discovery for human myocardium trained on triaxial shear and biaxial extension test for varying initial conditions. Discovered material parameters for simultaneous training with six shear and five biaxial tests with a regularization of *α* = 0.01 for five random parameter initializations; mean goodness of fit R^2^ and root mean squared error rms.

| network weights<br>$w_{1,\bullet}$ [-], $w_{2,\bullet}$ [kPa] | model term<br>[-] | random initialization | | | | |
| --- | --- | --- | --- | --- | --- | --- |
|  |  | #1 | #2 | #3 | #4 | #5 |
| $w_{1,7} \cdot w_{2,7}$ | $[I_2 - 3]^2$ | 5.162 | 5.162 | 5.171 | 5.153 | – |
| $w_{1,8}, w_{2,8}$ | $\exp([I_2 - 3]^2)$ | – | – | – | – | 1.390 3.482 |
| $w_{1,12}, w_{2,12}$ | $\exp([\bar{I}_{4f} - 1]^2)$ | 21.151, 0.081 | 21.174, 0.081 | 19.499, 0.093 | 21.146, 0.081 | 19.230, 0.098 |
| $w_{1,20}, w_{2,20}$ | $\exp([\bar{I}_{4n} - 1]^2)$ | 4.371, 0.315 | 4.373, 0.315 | 1.986, 0.784 | 4.371, 0.315 | 1.393, 1.062 |
| $w_{1,23}, w_{2,23}$ | $[I_{8fs}]^2$ | – | – | 0.335 | – | – |
| $w_{1,24}, w_{2,24}$ | $\exp([I_{8fs}]^2)$ | 0.508, 0.486 | 0.507 0.486 | – | 0.508, 0.486 | 0.586, 0.489 |
| $R^2$ | | 0.894 | 0.894 | 0.893 | 0.894 | 0.893 |
| rms |  | 0.426 | 0.426 | 0.428 | 0.426 | 0.430 |

### Our constitutive neural network specializes well to classical constitutive models

By constraining the majority of weights to zero and only training for a selective subset of weights [43], we can utilize our neural network to identify the parameters of popular classical constitutive models. We demonstrate this feature for three widely used orthotropic models for myocardial tissue: the Holzapfel Ogden model [33], the Guan model [26], and the generalized Holzapfel model [33]. The Holzapfel Ogden model is a *four-term model* [33] that features an exponential linear term in the first invariant *I*_1_, exponential quadratic terms in the fiber and sheet fourth invariants **Ī**_4f_ and **Ī**_4s_, and an exponential quadratic term in the fiber-sheet eighth invariant *I*_8fs_. We obtain the Holzapfel Ogden model by selectively training four sets of weights, *{w*_*º*,2_, *w*_*º*,12_, *w*_*º*,16_, *w*_*º*,24_*}*, while constraining all other weights to zero,

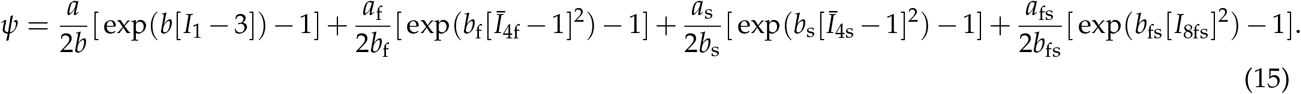

The Guan model is a slightly different *four-term model* [26] that features an exponential linear term in the first invariant *I*_1_, exponential quadratic terms in the fiber and normal fourth invariants **Ī**_4f_ and **Ī**_4n_, and an exponential quadratic term in the fiber-sheet eighth invariant *I*_8fs_. We obtain the special case of the Guan model by selectively training four sets of weights, { *w* _*º*,2_, *w* _*º*,12_, *w* _*º*,20_, *w* _*º*,24_}, while constraining all other weights to zero,

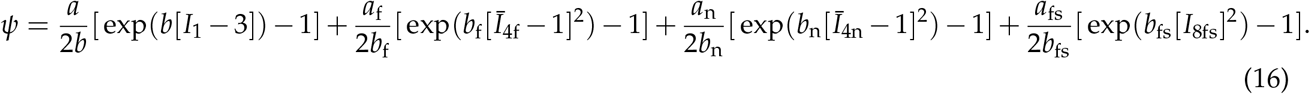

The generalized Holzapfel model is a *seven-term model* [33] that contains both previous models as special cases and features an exponential linear term in the first invariant *I*_1_, exponential quadratic terms all fourth invariants **Ī**_4f_, **Ī**_4s_, **Ī**_4n_, and an exponential quadratic term in all eighth invariant *I*_8fs_, *I*_8fn_, *I*_8sn_. We obtain the special case of the Guan model by selectively training seven sets of weights, *{w*_*º*,2_, *w*_*º*,12_, *w*_*º*,16_, *w*_*º*,20_, *w*_*º*,24_, *w*_*º*,28_, *w*_*º*,32_*}*, while constraining all other weights to zero,

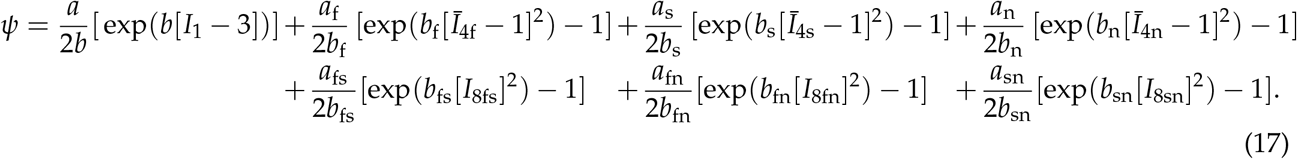

We emphasize that our network weights translate directly into physically meaningful constitutive parameters with well-defined physical units, namely the stiffness like parameters with the unit of kilopascals, *a* = 2*w*_1,2_*w*_2,2_, *a*_f_ = 2*w*_1,12_*w*_2,12_, *a*_s_ = 2*w*_1,16_*w*_2,16_, *a*_n_ = 2*w*_1,20_*w*_2,20_, *a*_fs_ = 2*w*_1,24_*w*_2,24_, *a*_fn_ = 2*w*_1,28_*w*_2,28_, *a*_sn_ = 2*w*_1,32_*w*_2,32_, and the unit-less nonlinearity parameters, *b* = *w*_1,2_, *b*_f_ = *w*_1,12_, *b*_s_ = *w*_1,16_, *b*_n_ = *w*_1,20_, *b*_fs_ = *w*_1,24_, *b*_fn_ = *w*_1,28_, *b*_sn_ = *w*_1,32_. To compare these three models against our discovered model and against each other, we constrain our network and train selectively for their non-zero weights. Figures 6 and 7 and show the stress-shear and stress-stretch plots of the Holzapfel Ogden and Guan models, and Table 4 summarizes the resulting network weights. Notably, the classical four-term Holzapfel Ogden model in Eq. (15) displays limitations when calibrated simultaneously for triaxial shear and biaxial extension; its stress plots in Figure 6 result in a mean goodness of fit as low as R^2^ = 0.788 and a root mean squared error of rms = 0.544. The four-term Guan model in Eq. (16) results in a visibly improved fit of the stress plots in Figure 7, with an improved mean goodness of fit of R^2^ = 0.867 and a root mean squared error of rms = 0.442. Naturally, the generalized seven-term Holzapfel model in Eq. (17) provides the most freedom to fit the data and results in an even better mean goodness of fit of R^2^ = 0.876 and a root mean squared error of rms = 0.440, yet at the cost of two additional terms and four additional parameters. Interestingly, our discovered three- and four-term models with a mean goodness of fit of R^2^ = 0.872 and R^2^ = 0.894 and only five and seven parameters both outperform the classical four-term Holzapfel Ogden and Guan models. Taken together, these results confirm that our constitutive neural network contains classical constitutive models as special cases and can successfully identify their parameters by constraining as large subset of weights to zero.

**Table 4:** Model specification for human myocardium fit for triaxial shear and biaxial extension tests. Identified material parameters for Holzapfel Ogden, Guan, and generalized Holzapfel models, from simultaneous training with six shear and five biaxial tests; mean goodness of fit R^2^ and root mean squared error rms.

| network weights<br>$w_{1,\bullet}$ [-], $w_{2,\bullet}$ [kPa] | model term<br>[-] | neural network special case | | |
| --- | --- | --- | --- | --- |
|  |  | Holzapfel Ogden Eq. (15) | Guan Eq. (16) | gen. Holzapfel Eq. (17) |
| $w_{1,2}, w_{2,2}$ | $\exp(I_1 - 3)$ | 3.949, 0.297 | 7.248, 0.054 | 5.457, 0.087 |
| $w_{1,12}, w_{2,12}$ | $\exp([\bar{I}_{4f} - 1]^2)$ | 14.389, 0.114 | 14.571, 0.154 | 23.701, 0.070 |
| $w_{1,16}, w_{2,16}$ | $\exp([\bar{I}_{4s} - 1]^2)$ | – | – | 20.067, 0.035 |
| $w_{1,20}, w_{2,20}$ | $\exp([\bar{I}_{4n} - 1]^2)$ | – | 10.929, 0.115 | 16.976, 0.060 |
| $w_{1,24}, w_{2,24}$ | $\exp([I_{8fs}]^2)$ | – | 4.959, 0.044 | 1.081, 0.271 |
| $w_{1,32}, w_{2,32}$ | $\exp([I_{8sn}]^2)$ | – | – | 11.842, 0.002 |
| $R^2$ | | 0.788 | 0.867 | 0.876 |
| rms |  | 0.554 | 0.442 | 0.420 |

**Figure 6:**
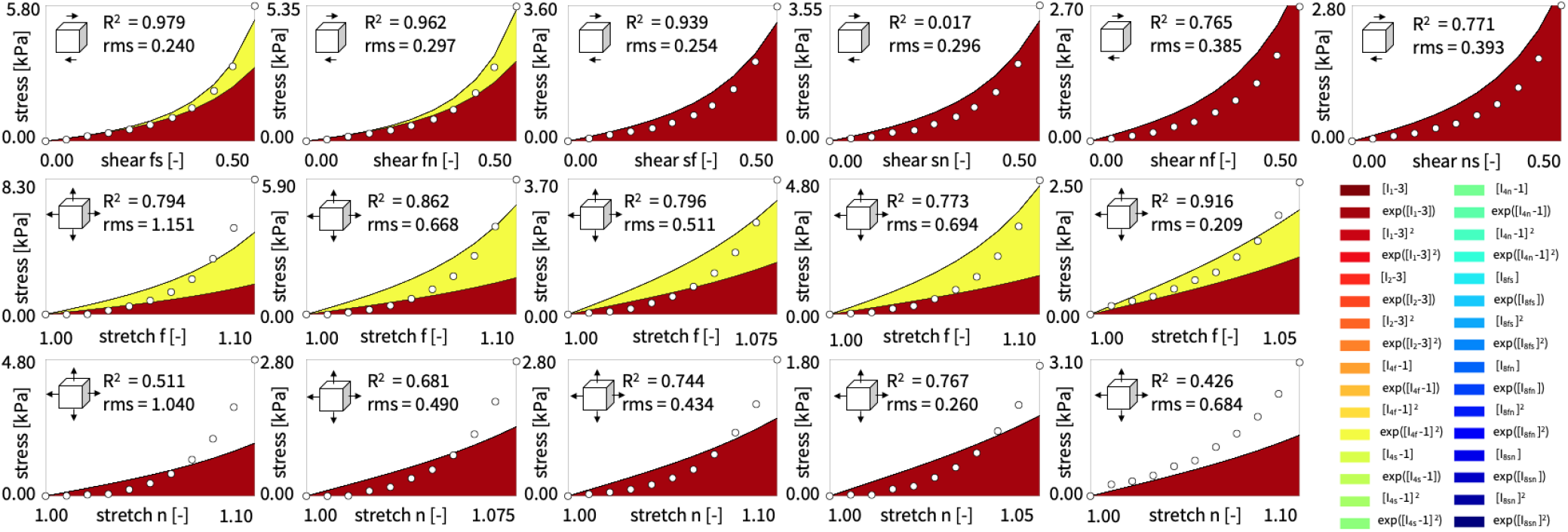
Parameter identification for Holzapfel Ogden model trained on triaxial shear and biaxial extension tests simultaneously. Cauchy stress components as functions of shear strain during triaxial shear, first row, and stretches during biaxial extension in fiber and normal directions, second and third rows, all used for training of the orthotropic, perfectly incompressible constitutive neural network with two hidden layers constrained to four nodes [33]. Dots illustrate the the experimental data [61]; color-coded areas highlight the four contributions to the discovered stress function.

**Figure 7:**
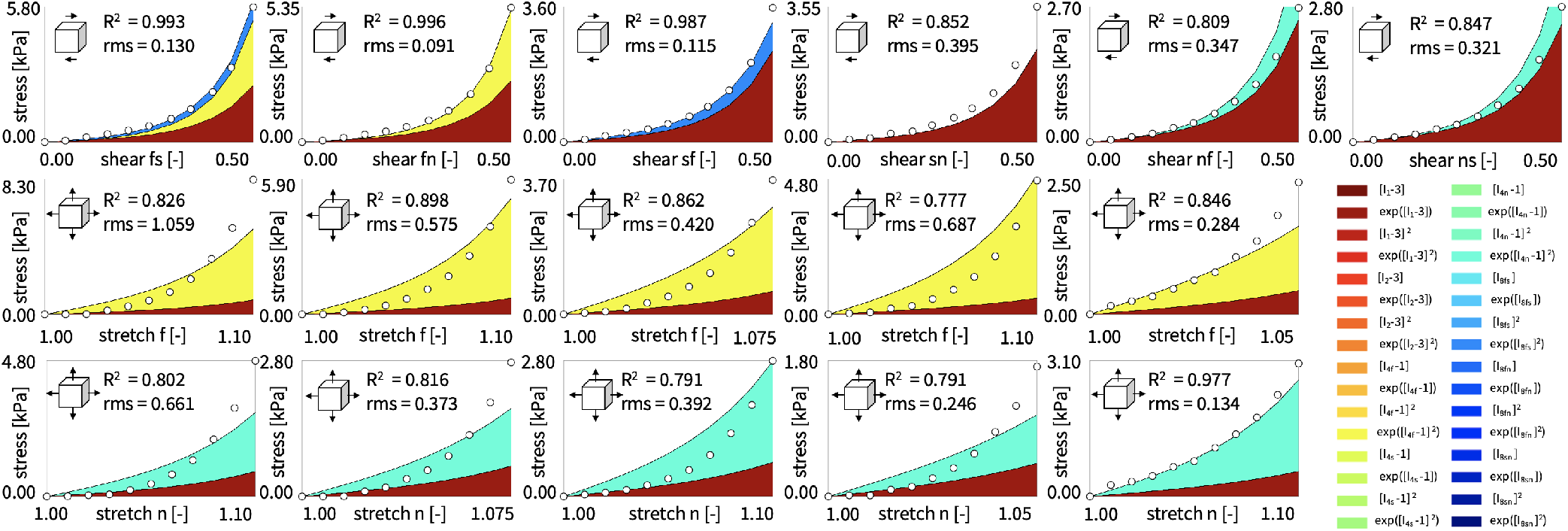
Parameter identification for Guan model trained on triaxial shear and biaxial extension tests simultaneously. Cauchy stress components as functions of shear strain during triaxial shear, first row, and stretches during biaxial extension in fiber and normal directions, second and third rows, all used for training of the orthotropic, perfectly incompressible constitutive neural network with two hidden layers constrained to four nodes [26]. Dots illustrate the the experimental data [61]; color-coded areas highlight the four contributions to the discovered stress function.

### Our discovered models generalize well from homogeneous tissue tests to real heart simulations

To explore whether our discovered models not only explain the behavior of myocardial tissue in the laboratory setting, but generalize to realistic human heart simulations, we predict the stress profiles across the left and right ventricular walls during diastolic filling. Specifically, we apply endocardial pressures of 8mmHg and 4mmHg in each ventricle, to mimic the healthy ventricular end diastolic pressure states. Figures 8 and 9 summarize the long-axis, short-axis, frontal, and top views of wall stress predictions for discovered model of different complexity: the four-term model, *ψ* = *w*_1,7_*w*_2,7_[*I*_2_ *−* 3]^2^ + *w*_2,12_[exp(*w*_1,12_[**Ī**_4f_ *−* 1]^2^) *−* 1] + *w*_2,20_[exp(*w*_1,20_[**Ī**_4n_ *−* 1]^2^) *−* 1] + *w*_2,24_[exp(*w*_1,24_[*I*_8fs_]^2^) *−* 1] for *α* = 0.01, the three-term model, *ψ* = *w*_1,7_*w*_2,7_[*I*_2_ *−* 3]^2^ + *w*_2,12_[exp(*w*_1,12_[**Ī**_4f_ *−* 1]^2^) *−* 1] + *w*_2,20_[exp(*w*_1,20_[**Ī**_4n_ *−* 1]^2^) *−* 1] for *α* = 0.1, and the one-term model *ψ* = *w*_2,6_[exp(*w*_1,6_[*I*_2_ *−* 3]) *−* 1] for *α* = 1.0, from left to right, with model parameters from Table 2. In this direct side-by-side comparison of the maximum principal stress profiles, we observe an excellent agreement for both *anisotropic* models, the *α* = 0.01 four-term model and the *α* = 0.1 three-term model. Strikingly, the *isotropic* model, the *α* = 1.0 one-term model, performs almost identically, with only minor local differences in the form of reduced stresses along the septum and across the left endocardial wall. We can easily understand these differences when comparing the performance of the discovered *α* = 0.01, *α* = 0.1, and *α* = 1.0 models during the homogeneous laboratory testing in Figure 5, where the *α* = 1.0 model overestimates the stresses in the low-stretch regime, but underestimates the stresses in the high-stretch regime. In the real heart simulations of Figure 8, this high-stretch regime is located along the left endocardial wall, where the differences between the models are most pronounced. In the frontal and top views of Figure 9, these differences are barely visible and most characteristic features are captured equally by all three models, including the isotropic model with only a single term. Taken together, our discovered models generalize well from homogeneous tissue tests to real heart simulations, with an unexpectedly accurate performance of the simplest isotropic model with a single exponential second invariant term. Figures 10 and 11 summarize the long-axis, short-axis, frontal, and top views of wall stress predictions for our discovered four term model in Eq. (13) compared to popular existing models, the Holzapfel Ogden model in Eq. (15), the Guan model from Eq. (16), and the generalized Holzapfel model from Eq. (17), with model parameters from Table 4. Interestingly, our side-to-side comparison showcases remarkably similar maximum wall stresses across all four models. In the short-axis and longaxis views of Figure 10, we observe small quantitative differences between the maximum principal wall stresses across the endocardial left ventricular free wall, with larger values for our discovered model and the Guan model and smaller values for the Holzapfel Ogden model. Again, we can explain these differences by comparing the performance of these three models during the homogeneous laboratory testing in Figures 5, 6, and 7, where the Holzapfel Ogden model underestimates the stresses in the high-stretch regime, while our discovered model and the Guan model approximate these stresses more accurately.

**Figure 8:**
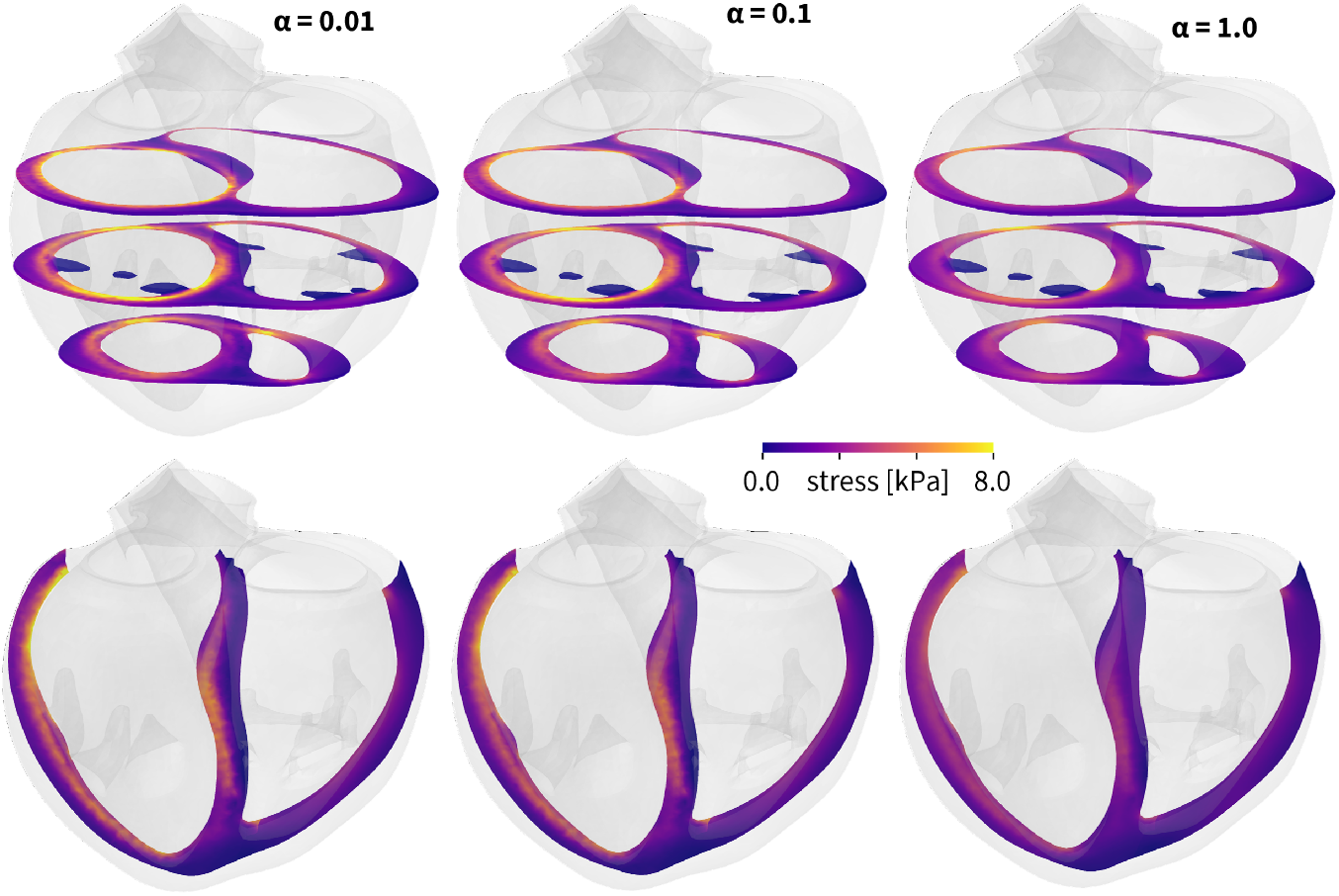
Stress profiles across the human heart, in short-axis and long-axis views, predicted by our discovered material models. Maximum principal stresses generated by a healthy left and right ventricular end-diastolic pressure of 8mmHg and 4mmHg. Predictions with three different models for varying regularization levels, discovered four-term model for *α* = 0.01, three-term model for *α* = 0.1, and one-term model for *α* = 1.0 with parameters from Table 2.

**Figure 9:**
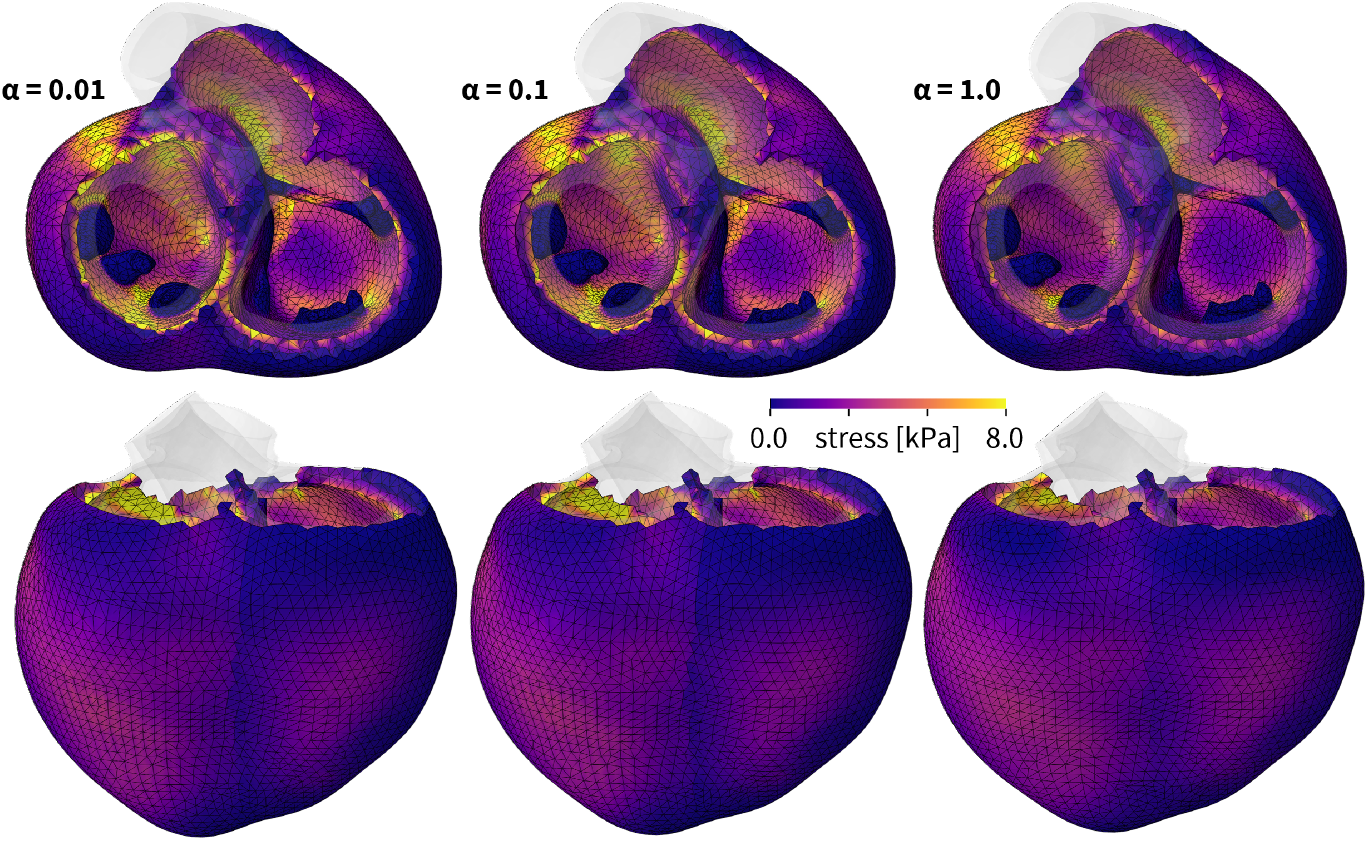
Stress profiles across the human heart, in a frontal and top views, predicted by our discovered material models. Maximum principal stresses generated by a healthy left and right ventricular end-diastolic pressure of 8mmHg and 4mmHg. Predictions with three different models for varying regularization levels, discovered four-term model for *α* = 0.01, three-term model for *α* = 0.1, and one-term model for *α* = 1.0 with parameters from Table 2.

**Figure 10:**
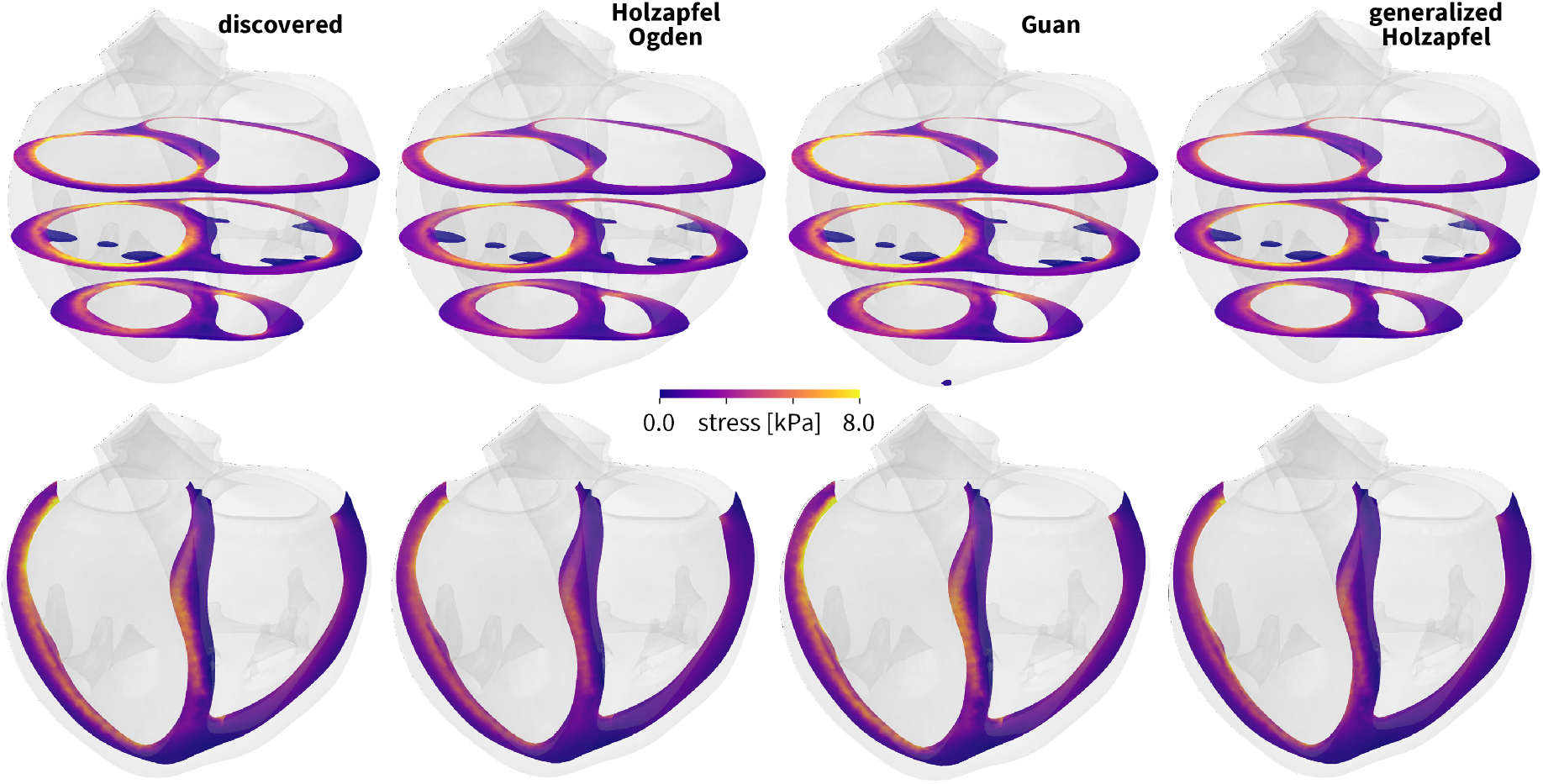
Stress profiles across the human heart, in short-axis and long-axis views, predicted by our alternative discovered material models. Maximum principal stresses generated by a healthy left and right ventricular end-diastolic pressure of 8mmHg and 4mmHg. Predictions for four different models, our discovered four-term model for *α* = 0.01 with parameters from Table 2, and the Holzapfel Ogden, Guan, and generalized Holzapfel models with parameters from Table 4.

**Figure 11:**
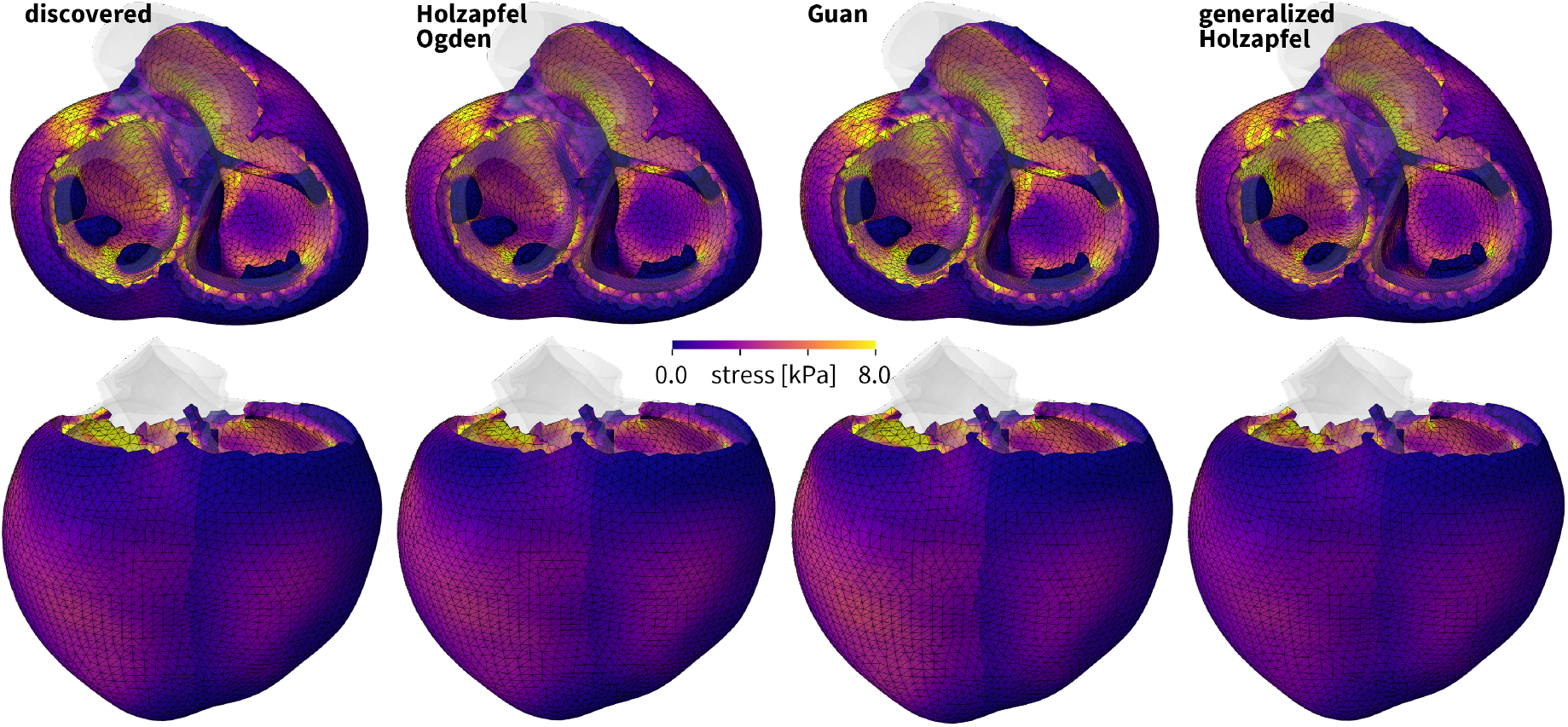
Stress profiles across the human heart, in a frontal and top views, predicted by our alternative discovered material models. Maximum principal stresses generated by a healthy left and right ventricular end-diastolic pressure of 8mmHg and 4mmHg. Predictions for four different models, our discovered four-term model for *α* = 0.01 with parameters from Table 2, and the Holzapfel Ogden, Guan, and generalized Holzapfel models with parameters from Table 4.

This is in line with the lowest goodness of fit for the Holzapfel Ogden model of R^2^ = 0.788, compared to our discovered model with R^2^ = 0.894, the Guan model with R^2^ = 0.867, and the generalized Holzapfel model with R^2^ = 0.876. Additionally, our diastolic hemodynamic loading enforces deformation and stress states that surpass the homogeneous tissue testing protocols of the triaxial shear and biaxial extension training data, which creates local regions of extrapolation beyond the initial training regime. Taken together, while our discovered four-parameter model best explains the laboratory experiments of triaxial shear and biaxial extension, all four models translate well into a single universal material subroutine and predict fairly similar stress profiles across the human heart.

## 4 Discussion

The objective of this work was to discover the model and parameters that best describe the mechanical behavior of human cardiac tissue. Towards this goal, we adopt the paradigm of constitutive neural networks, supplemented by *L*_*p*_-regularization. We explore and discuss the most important features and challenges of model discovery, with a view towards selecting appropriate training data, sparsifying the discovered model, comparing the model to popular existing models, and generalizing it from homogeneous training data to heterogenous real heart simulations.

### Our constitutive neural network discovers sparse and interpretable models to explain human cardiac tissue

Our constitutive neural network in Figure 1 features 32 individual terms, 8 isotropic and 24 anisotropic. This results in 2^32^ possible combinations of terms, a total of more than 4 billion models, represented through 64 network weights. Of these 64, all odd weights of the first layer are redundant, and we can set them equal to one to reduce the total number of trainable weights to 48. We first train the network with three different data sets, triaxial shear, biaxial extension, and shear and extension combined [61], initially without any regularization. From Figures 2, 3, and 4, we conclude that the network trains well for all three data sets, and successfully discovers models to explain the training data. However, the discovered models are not sparse; they contain a large number of terms and a large set of non-zero parameters [63]. For example, in Figure 3, top row, we observe that, although we only use biaxial extension data for training, the network activates terms and parameters that are not associated with any of the five biaxial extension tests: Although training with biaxial extension in the fn-plane does not provide any information about the shear behavior in the fs- and sn-planes, the weights related to the eighth invariants *I*_8fs_ and *I*_8sn_ are non-zero and contribute to the shear response. Strikingly, this issue resolves itself naturally when training with triaxial shear and biaxial extension combined. For training with both data sets simultaneously, three quarter of all weights train to zero, even in the complete absence of regularization.

Figure 4 highlights the remaining non-zero terms and Table 2, third column, summarizes their parameter values. However, these unregularized models are still fairly complex, sensitive to noise, and computationally expensive [9, 42]. This includes both, convergence during training, as we conclude from the required number of epochs in Table 2, and performance during simulations. To induce sparsity, we supplement the loss function in Eq. (8) with *L*_1_-regularization or lasso [48, 67]. Table 2 confirms that increasing the regularization level *α* from zero to one induces sparsity by gradually dropping the weights that have the smallest influence on the loss function. Figure 5 visualizes this reduction in model complexity, from left to right, associated with a decreasing number of colors, from eight to one, but also emphasizes the associated reduction of the goodness of fit. Taken together, we conclude that our *L*_1_-regularized constitutive neural network can reliably discover sparse and interpretable models and physically meaningful parameters to explain the complex behavior of human cardiac tissue.

### Our constitutive neural network is a generalization of popular constitutive models

By constraining the majority of weights to zero and only training for a selective subset of weights [43], we can utilize our neural network to identify the parameters of popular classical constitutive models. As a matter of fact, our neural network in Figure 1 is a generalization of previous invariant-based neural networks for isotropic materials [43] and for transversely isotropic materials [44] and naturally captures all their features as special cases. As such, we can reduce it to represent popular *isotropic* models including the neo Hooke [68], Blatz Ko [8], Mooney Rivlin [49, 58], or Demiray [12] models, as well as *transversely isotropic* models including the Lanir [39], Weiss [71], Groves [25], or Holzapfel [31] models. Figures 6 and 7 and Table 4 confirm that we can also reduce our neural network to represent popular *orthotropic* models including the Holzapfel Ogden [33], Guan [26], and generalized Holzapfel [33] models. Interestingly, the objective of the Guan model [26] was to systematically reduce the 14-parameter generalized Holzapfel model [33] using the Akaike information criterion, for the same experimental data of human myocardium that we used in this study [61]. The Akaike information criterion rewards the goodness of fit and penalizes the number of parameters with the goal to induce sparsity and prevent overfitting [2]. This reduces the generalized Holzapfel model with seven terms and 14 parameters in Eq. (17) to the Guan model with four terms and eight parameters in Eq. (16). Strikingly, when using *L*_1_-regularization with a penalty parameter *α* = 0.01, we discover exactly the same three anisotropic terms as the Guan model [26]: exponential quadratic terms in the invariants **Ī**_4f_, **Ī**_4n_, *I*_8fs_, associated with terms 12, 20, 24 of our constitutive neural network, colorcoded in yellow, turquoise, and blue. Yet, our discovered four-term model with a mean goodness of fit of *R*^2^ = 0.894 and an error of 0.426 outperforms the Guan model with a mean goodness of fit of *R*^2^ = 0.867 and an error of 0.442. Notably, while both models share these *same anisotropic terms*, they feature a *different isotropic term*, the classical exponential linear *I*_1_ term in the Guan model [26] and the quadratic *I*_2_ term in our discovered model.

### Our constitutive neural network consistently discovers second-invariant models

For decades, the gold standard in constitutive modeling has been to *first* select a constitutive model and *then* fit the model to data [5, 6, 17, 32, 60, 69]. Attempts to improve the goodness of fit have slightly adjusted the terms of the model, and gradually modified or replaced individual terms [11]. Admittedly, this has probably been the *only* way to improve constitutive models, simply because of the extreme non-linearity associated with this problem, its non-convexity, its multiple local minima, and the shear computational complexity associated with finding a good constitutive model. Now, with the massive advancement of computational power and the development of fast and efficient solvers, a unique opportunity presents itself to *simultaneously* discover both the best model and the best parameters to explain experimental data [42]. This opens doors to probe a huge variety of common functional building blocks [43], in our case 32, and automatically select the best combination of terms, in our case out of more than 4 billion possible combinations. Traditionally, cardiac tissue models have a priori postulated that the isotropic behavior is best described by the *first invariant* of the strain, and ignored the *second invariant* [12, 20, 24, 33, 35]. This is in line with dozens of constitutive models for other biological systems and natural and man-made soft matter [22, 32]. Intriguingly, our constitutive neural network allows us to probe both invariants simultaneously, not only in their linear or quadratic forms, but also embedded in exponentials, not only in isolation, but also in combination with other anisotropic terms [44]. This would have been unthinkable a decade ago! The models we discover using this approach display as striking yet consistent trend: All discovered models feature the second invariant instead of the first. Importantly, this observation is not exclusive to sparsification, it reflects a universal trend that is present, even in the complete absence of regularization. Table 2 confirms that both the non-regularized network in column three and the regularized networks in columns four to eight only discover models in terms of the second invariant *I*_2_. Visually, we can easily confirm this selective activation in the color-coded stress terms in Figure 4, which prominently display orange-type colors associated with the second invariant. The remarkable dominance of the second invariant is in stark contrast with the popularity of classical models that only feature the first invariant [10, 12, 31, 33, 39, 68], but consistent with recent models for soft biological tissues [8, 25, 34, 49, 58, 71]. Notably, for triaxial shear testing, the first and second invariants are identical, *I*_1_ = *I*_2_ = *γ*^2^ + 3. This implies that their differentiation is meaningless if the model is trained on triaxial shear experiments alone. This explains why the classical Holzapfel Ogden model, fit only to six *triaxial shear* experiments [33], performs exceptionally well, although it only uses the first invariant. When using *biaxial extension* experiments, the first and second invariant are no longer identical and the model consistently favors the second invariant over the first. Taken together, in agreement with previous observations [43, 56], we find that the second invariant is better suited to capture the isotropic response of biological tissues [34] and describes the experimental data more accurately than the first.

### Our constitutive neural network consistently discovers exponential quadratic terms

Prior to the now widely used invariant-based Holzapfel-type models [31], the common standard to model the anisotropic behavior of arterial and cardiac tissues were strain-based Fung-type models [21] that simply embedded a combination of strains into an exponential free energy function. The fundamental difference between both families of models is that Fung-type models draw motivation from the *macrostructural* orientation encoded in radial, circumferential, and longitudinal directions [22], while Holzapfel-type models are inspired by the *microstructural* architecture encoded in fiber, sheet, and normal orientations [32]. Our constitutive neural network models anisotropy using an invariant-based microstructural approach [44]. Yet, rather than using a limited number of invariants, our network offers the full set of three fourth invariants *I*_4f_, *I*_4s_, *I*_4n_ and three eights invariants *I*_8fs_, *I*_8fn_, *I*_8sn_. And rather than using a specific functional form, our network offers linear, exponential linear, quadratic, and exponential quadratic activation functions (º), (exp(º)), (º)^2^, (exp((º)^2^)) to each of these invariants. Notably, this results in a total of (6 *×* 4)^2^, more than 16 million, of possible combinations of anisotropic terms. Strikingly, of all these combinations, our network consistently favors models with *exponential quadratic terms*. Tables 2 and 3 suggest that the dominance of these exponential quadratic terms is independent of the regularization level and independent of the initial conditions. This observation stands in contrast to the popularity of earlier anisotropic models including the linear fourth invariant Lanir model [39] and the exponential linear Demiray [12], Weiss [71], and Groves [25] models, but is in line with the massive popularity of the exponential quadratic Holzapfel model for both arteries [31] and cardiac tissue [33]. Taken together, our automated model discovery consistently discovers and confirms the widely used exponential quadratic terms that have been introduced more than two decades ago to model the strain-stiffening behavior of collagen fibers in soft biological tissues.

### Limitations

Our results demonstrate that we can successfully adopt constitutive neural networks to discover a model and a set of physically meaningful parameters that best describe the behavior of human cardiac tissue. However, we encountered a few limitations that point towards future investigations: First, while we have discovered the best model to explain the *available data*, the data themselves might be biased towards probing more in the fiber and normal plane, which could explain why we have prominently discovered *I*_4f_ and *I*_4n_ terms instead of *I*_4s_. Second, we have assumed that cardiac tissue is *perfectly incompressible*, a limitation that we could address by adding the third invariant and learning the bulk modulus as additional network weight, provided we have sufficient experimental data. Third, we have assumed that the tissue is *hyperelastic*, a limitation that we could address by incorporating an inelastic potential, for example, to account for time-dependent viscoelastic effects. Fourth, our network architecture in Figure 1 is limited to models with *decoupled invariants*, which we could address by using a more densely connected architecture in which some or all nodes between the first and second hidden layers are interconnected.

Fifth, at times, our method is sensitive to its *initialization*, which we view as strength rather than weakness, since it allows us to explore alternative models with different combinations of terms. Sixth, while we currently assume that we know the fiber, sheet, and normal orientations, we could introduces these *microstructural features* as trainable parameters and discover them alongside the macroscopic model parameters. To address any of these limitations, it would be tremendously useful to acquire additional data, ideally from tension and compression tests, in isolation and in combination with shear, for quasi-static loading, and for loading at different rates.

## Conclusion

For more than five decades, scientists have been trying to develop constitutive models for the heart. While most models work well for individual tests such as tension, compression, or shear, each model has its own limitations when fit simultaneously to a combination of tests. Here, instead of a priori selecting a specific model and fitting its parameters to data, we simultaneously discover the best model and parameters using incompressible orthotropic constitutive neural networks. We train our network using six triaxial shear and five biaxial extension tests and sparsify the resulting model using *L*_1_-regularization. Our results suggest that an accurate material model for cardiac tissue should at least include one isotropic and two or three anisotropic terms. Strikingly, instead of a linear first invariant term, the network consistently discovers a quadratic second invariant term to best represent the isotropic response. Notably, to model the anisotropic response, the network discovers two exponential quadratic fourth invariant terms that resemble the classical Holzapfel format. Importantly, all our discovered weights have a clear physical interpretation and translate into stiffness-like and nonlinearity parameters. Finally, we embedded all discovered models into a finite element simulation to predict the stress profiles across the human heart during diastolic filling, and compared them against other popular cardiac models. Our results suggest that constitutive neural networks can successfully discover interpretable and generalizable model and parameters to accurately simulate and predict deformations and stresses in the human heart in real life situations. We anticipate that our new four-term model for cardiac tissue will have broad applications in biomedical device design, medical diagnostics, and management of cardiovascular disease.

## Data availability

Our source code, data, and examples are available at https://github.com/LivingMatterLab/CANN.

## Acknowledgments

We thank Jiang Yao, Juan A. Hurtado, and the Living Heart Project team from Dassault Systèmes for their helpful discussions and suggestions. This work is part of the I2 ♥ I2 movement that started in Palo Alto in February 2024. It was supported by the DFG Grant 496647562 to Denisa Martonová, by the NWO Veni Talent Award 20058 to Mathias Peirlinck, and by the NSF CMMI Award 2320933 Automated Model Discovery for Soft Matter to Ellen Kuhl.

## Notes

### Competing Interest Statement

The authors have declared no competing interest.

